# Phylogenetic network analysis revealed the recombinant origin of the SARS-CoV-2 VOC202012/01 (B.1.1.7) variant first discovered in U.K.

**DOI:** 10.1101/2021.06.24.449840

**Authors:** Xianfa Xie, Teash-Juan Lewis, Nikoli Green, Zhenping Wang

## Abstract

The emergence of new variants of the SARS-CoV-2 virus poses serious problems to the control of the current COVID-19 pandemic. Understanding how the variants originate is critical for effective control of the spread of the virus and the global pandemic. The study of the virus evolution so far has been dominated by phylogenetic tree analysis, which however is inappropriate for a few important reasons. Here we used phylogenetic network approach to study the origin of the VOC202012/01 (Alpha) or so-called UK variant (PANGO Lineage B.1.1.7). The multiple network analyses using different methods consistently revealed that the VOC202012/01 variant was a result of recombination, in contrast to the common assumption that the variant evolved from step-wise mutations in a linear order. The study provides an example for the power and application of phylogenetic network analysis in studying virus evolution, which can be applied to study the evolutionary processes leading to the emergence of other variants of the SARS-CoV-2 virus as well as many other viruses.

**Significance:** The emergence of new variants of the SARS-CoV-2 virus, including the Alpha variant first found in U.K., poses serious challenges to the control of the current COVID-19 pandemic. Understanding how new variant originated is paramount to end the pandemic as effectively and quickly as possible. The dominant phylogenetic tree approach to study virus evolution has been inadequate and even misleading. Here we used a phylogenetic network approach to study the origin of the VOC202012/01 (Alpha) variant which was first reported in U.K. last year but has soon spread into many other countries, leading to dramatic increase in infection and death. Multiple analyses consistently revealed that the variant originated through recombination of pre-existing virus strains, highlighting an important but largely ignored mechanism in the evolution of the SARS-CoV-2 virus so far.

---

The global COVID-19 pandemic has caused over 146 millions of cases and over 3 millions of deaths around the globe by the time this manuscript is written (John Hopkins University Coronavirus Resource Center 2021). The causing virus, SARS-CoV-2, can be spread mostly through the respiratory system but may also be spread through fecal-oral or fecal aerosol transmissions (Jiao *et al*. 2021, Meng and Liang 2021). Just like many other types of viruses, the SARS-CoV-2 virus has kept evolving into new variants. The so-named “UK variant”, which should be more properly called the Variant of Concern (VOC) 202012/01 or Alpha variant as it is currently named, was first reported in the United Kingdom in December 2020 but has quickly spread into many other countries across different continents, leading to a surge of COVID-19 cases and deaths. Published study showed that the variant, which is classified as PANGO lineage B.1.1.7, has much higher transmissibility over other strains that had been in circulation in human populations (Davies *et al*. 2021). The emergence and rapid spread of the U.K. variant and other variants has serious implications for the control of COVID-19, including both detection and the effectiveness of the vaccines (McCarthy *et al*. 2021). Therefore, understanding how the U.K. variant had emerged at the first place could provide critical information on the processes producing new SARS-CoV-2 variants and help inform develop better strategies for pandemic control.

So far, almost all the studies of the evolution and spread of the SARS-CoV-2 virus have been based on phylogenetic tree analysis, including the representations at the GISAID website (Elbe and Buckland-Merrett 2017), which has been one of the most important databases for the genomic sequences of SARS-CoV-2, the widely popular Nextstrain website (Hadfield *et al*. 2018), and numerous publications on the subject. While phylogenetic tree analysis can be relatively easily conducted with well-established methods and produce easier to interpret results, the approach unfortunately is not appropriate for the evolutionary study of the SARS-CoV-2 viruses for the following reasons. First, phylogenetic tree construction forces all existing sequences to be at the tips of the tree and artificially prevents any existing sequence to be an ancestor to some other sequences. However, given the very short evolutionary history of the different strains of the SARS-CoV-2 viruses, some ancestral sequences which gave rise to other sequences may still be in circulation in the human population and might have been sampled by researchers. Second, bifurcating phylogenetic trees do not allow the possibility for one ancestral sequence to give rise to more than two descendant lineages, the latter of which might well have been the possibility in the evolution of the SARS-CoV-2 viruses. Third and most importantly, phylogenetic tree construction assumes no recombination between or among any ancestral sequences. However, it has been shown that a variety of RNA viruses could recombine through template switching to form new sequences, including HIV (Shriner *et al*. 2004, Neher and Leitner 2010, Simon-Loriere *et al*. 2010), polioviruses (Savolainen-Kopra and Blomqvist 2010), bromovirus (Urbanowicz *et al*. 2005), influenza virus (Lindstrom *et al*. 2004), the Western equine encephalitis virus (Weaver 2006), and coronavirus (Jackwood *et al*. 2010). For the above reasons, phylogenetic network analysis would be a much more appropriate approach to study the evolution of the SARS-CoV-2 viruses.

However, so far there have been very few published network analysis of the SARS-CoV-2 viruses with only one (Forster *et al*. 2020) having received much attention but also great criticisms. Some criticized that the result in the published study could be misinterpreted regarding the origin of the SARS-CoV-2 virus (Chookajorn 2020). Similarly, Mavian *et al*. (2020) and Sanchez-Pacheco *et al*. (2020) argued that the sampling bias and incorrect rooting made the result of the network analysis in Forster *et al*. 2020 unreliable.

While the above published phylogenetic network analysis of some of the earliest sampled SARS-CoV-2 virus genomes were criticized for different reasons, the phylogenetic network approach itself should not be discarded. On the contrary, for the reasons mentioned above, phylogenetic network approach would be much more appropriate to study the evolution of the SARS-CoV-2 viruses than the phylogenetic tree approach. Here we employed different network analysis methods to study the origin of the VOC202012/01 or so-called “UK Variant” and found that it very likely had originated from recombination.

The VOC202012/01 (PANGO Lineage B.1.1.7) was first reported in the media in December 2020 though the first detection of the variant in southeast England could be traced back to September 2020. Since then, it had become the dominant lineage in the United Kingdom and spread to more than 114 countries worldwide (Davies *et al*. 2020). The new variant has been shown to have a 43-90% higher reproduction number (R) than pre-existing variants. It was first documented to contain 17 unique amino acid changes compared to the reference genome of the SARS-CoV-2 virus, including amino acid substitutions and deletions, with consecutive deletions being considered as a single evolutionary change. The variant was first reported in the GR clade of GISAID but later reported in the GH, GV and other GISAID clades as well. As the GISAID clades were classified based on linear phylogenetic tree analysis, the existence of the VOC202012/01 variant in multiple clades was the first indication of the existence of recombination in the evolution of the variant, as the alternative explanation, *i*.*e*., homoplasies with exactly the same 17 amino acid changes at the same locations were the result of independent mutations in different clades, was almost impossible in probability.

However, most researchers have assumed that the VOC202012/01 variant itself had originated through a step-wise mutational process. But a large-scale phylogenetic tree analysis of the virus sequences in U.K. (Lauring and Hodcroft 2021) failed to demonstrate step-wise mutations leading to the occurrence of the VOC202012/01 variant. Instead, the variant sequences could only be connected to the other sequences by a very long branch, which is another strong indicator that the sequences could well be recombinants (Schierup and Hein 2000). Here we examine in details this alternative hypothesis, *i.e.*, the VOC202012/01 variant might have originated through recombination of existing variant sequences, a hypothesis that has been largely ignored by the researchers on the SARS-CoV-2 viruses.

Using a dataset including the earliest collected VOC202012/01 variant genome sequence in U.K., the reference genome sequence, and others sequences containing individual or multiple amino acid changes found in the VOC202012/01 variant, we first constructed phylogenetic trees using three different methods: maximum likelihood, neighbor-joining, and maximum parsimony. First, trees constructed using the three different methods show different relationship between the VOC202012/01 variant sequence and the other sequences except for the most closely related sequence. While the three different methods are expected to show largely consistent phylogenetic tree topologies for clearly defined evolutionary history. Secondly, trees constructed using the three methods consistently showed very low bootstrap values in the phylogenetic relationship between the VOC202012/01 variant sequence and the other sequences. Both results indicate the uncertainty of the location of the VOC202012/01 variant sequence in phylogenetic tree analysis and thus its evolutionary relationship with other sequences in a tree framework, a strong indicator of recombination for its origin.

To further explore the possibility of the recombinant origin of the VOC202012/01 variant, we conducted network analysis using two different packages. The analysis using PopART (Leigh and Bryant 2005) clearly showed that the first VOC202012/01 variant sequence in U.K. was the result of recombination of two different lineages. In contrast, the reference genome sequence, which represents one of the earliest collected samples of the SARS-CoV-2 virus, was shown to be at the opposite end of the network, consistent with the expected evolutionary history of the virus.

To verify this result, we analyzed the same dataset using another commonly used package SplitsTree4 (Huson and Bryant 2006). The result (Figure 2) also showed that the first documented VOC202012/01 variant sequence in U.K. was a recombinant from two immediate parental sequences, while each of the latter resulted from recombination at other levels. Similar to the results from the PopArt network analysis, the SARS-CoV-2 viruses reference genome is also shown to be very close to the root of the network, while the first VOC202012/01 variant sequence in U.K. is found at the opposite end of the network, both of which are consistent with the expected evolutionary history of the virus.

**Figure 1.**
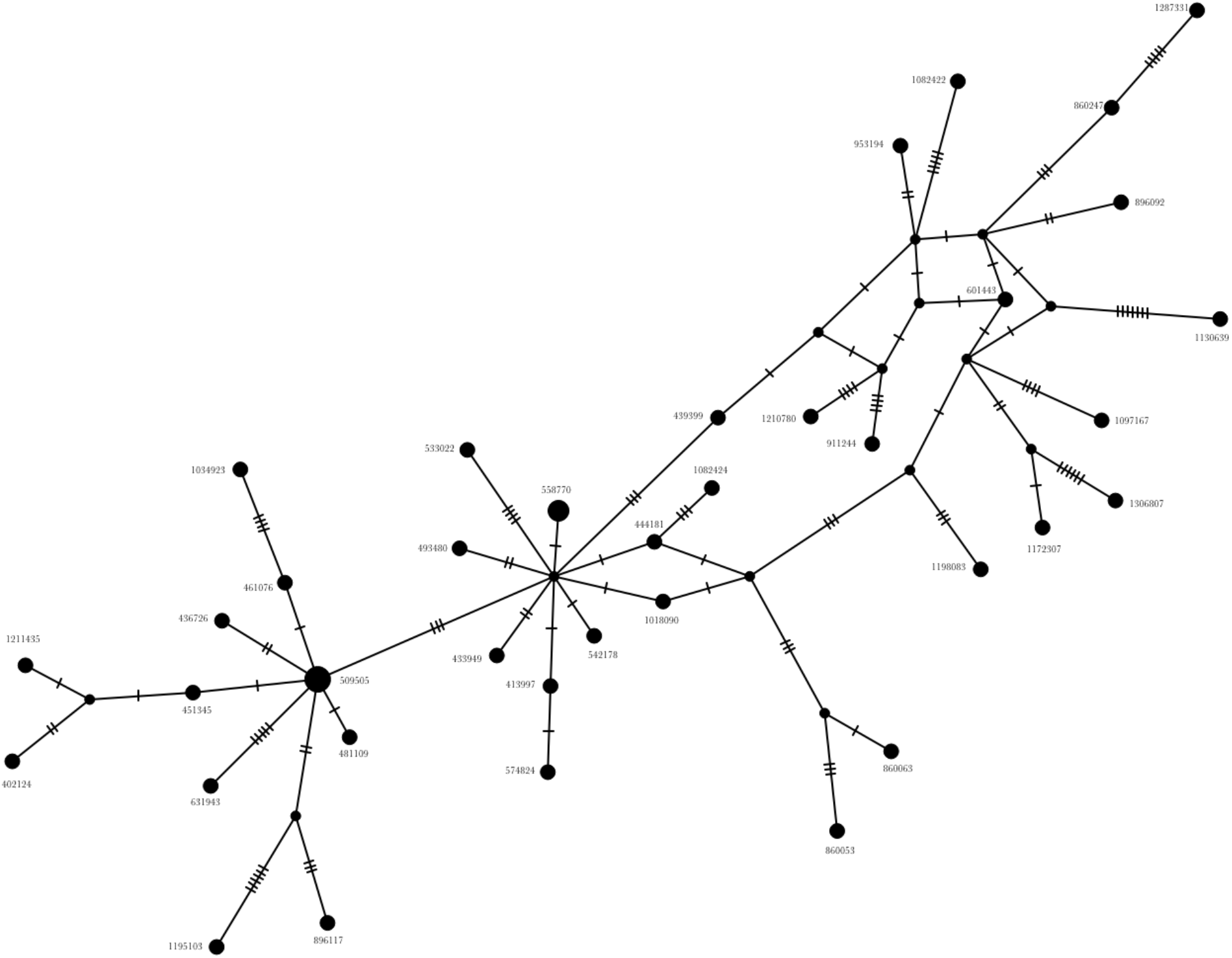
Network analysis with the PopART program. Note the reference genome sequence (EPI_ISL_402124) at one end of the network while the VOC202012/01 variant sequence (EPI_ISL_601443) at the other end of the network with possible recombinant origin.

**Figure 2.**
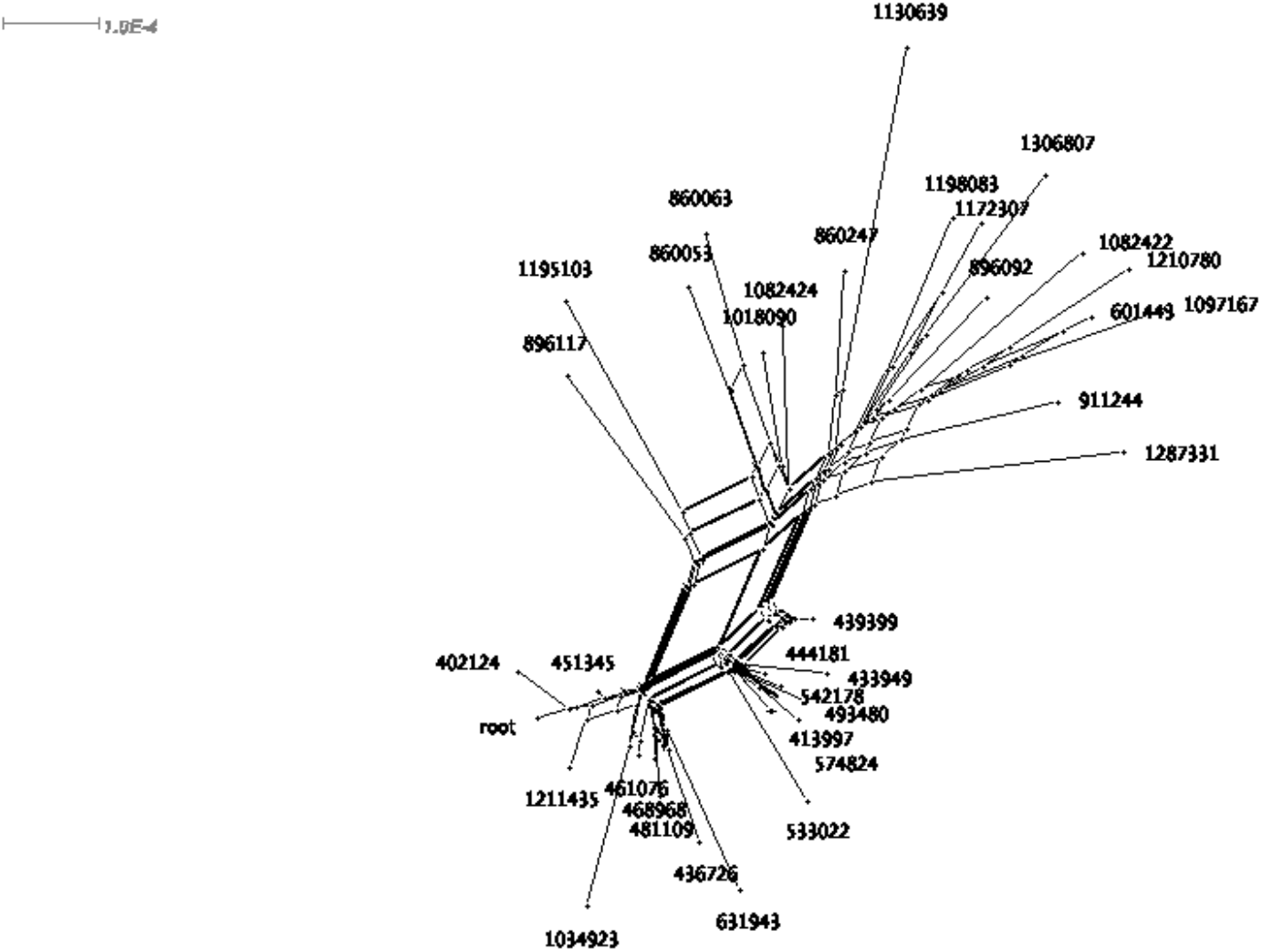
Network analysis with the SplitsTree4 program. The network is rooted with the reference genome sequence (EPI_ISL_402124) close to the root, as expected, and the VOC202012/01 variant sequence (EPI_ISL_601443) at the other end of the network with recombinant origin.

To test whether the recombinant origin of the first VOC202012/01 variant sequence in U.K. as shown in the above network analyses was coincidental, we added another VOC202012/01 variant sequence to the dataset and ran the above network analyses again. As shown in the supplemental Figures 2 and 3, the essential topologies in both analyses did not change at all except the newly added VOC202012/01 variant sequence was found to be derived from the first VOC202012/01 variant sequence in U.K, which is consistent with expectation.

In conclusion, our analyses demonstrated that the VOC202012/01 variant first identified in U.K. might have been the result of recombination among existing variants. This is a process that has been largely ignored for the study of the SARS-CoV-2 virus so far. However, as has been demonstrated through empirical research, recombination is common for many RNA viruses and can be of major evolutionary significance (Shriner *et al.* 2004, Neher and Leitner 2010, Simon-Loriere *et al.* 2010, Savolainen-Kopra and Blomqvist 2010, Urbanowicz *et al.* 2005, Lindstrom *et al.* 2004, Weaver 2006, Jackwood *et al.* 2010). It can happen through template switching during replication when different viruses co-infecting the same host (Simon-Loriere and Holmes 2011). A detailed study of the intra-host variation of SARS-CoV-2 during the early epidemic in the U.K. suggested that 1-2% of the samples could have been co-infected (Lythgoe *et al.* 2021), which provided many opportunities for new variants to emerge through recombination. However, the VOC202012/01 variant may not be the only variant that originated through recombination. It is reasonable to suspect that some other variants that have recently emerged, including those in South Africa, Brazil, and India, might have been formed through recombination of existing variants as well. Our study also demonstrated the power of phylogenetic network analysis and its competitive advantages over phylogenetic tree analysis in situations like the evolutionary study of the SARS-CoV-2 viruses. And this approach is currently being used to study the origin of some other variants, including the ones mentioned above. The recombinant origin of the VOC202012/01 variant underlies the importance of both local and global control of the pandemic as rapid as possible, as attenuation of the pandemic in any population may facilitate the emergence of new variants that may start new waves of infection and make it impossible to completely eliminate the SARS-CoV-2 virus in circulation in human population, though new vaccine production methods (Maeda *et al*. 2021) may offer hope for world-wide production of the vaccines against the recently emerged and still emerging variants of the SARS-CoV-2 virus.

## Materials and Methods

All the genomic sequences used in the analysis, including the reference genome sequence, were from the GISAID website (https://www.gisaid.org). The sequence with the earliest sample collection date in U.K. for the VOC202012/01 variant was identified to be EPI_ISL_601443, which was collected in England on September 20, 2020. Multiple VOC202012/01 variant sequences from different GISAID clades were compared to identify the shared amino acid changes in the VOC202012/01 variant. Subsequently, sequences containing individual or combinations of the amino acid changes found in the VOC202012/01 variant were identified and the sequences with the earliest sample collection date and fewest additional amino acid changes were selected whenever possible. It should be noted that the earliest sample collection date does not necessarily reflect the date the virus haplotype first emerged in human population because of inadequate sampling and the lag between emergence time and sample collection time, which is the reason some sequences collected after September 20, 2020 were also used for this study. And because the close proximity of U.K. to other countries in the Europe and the lack of complete shutdown among the countries during the pandemic, sequences from other European countries were also used as long as they met the above criteria. In all the searches in the GIDAID database, however, the following filters were applied: complete, high coverage, low coverage excluded, and collection date complete.

The first VOC202012/01 variant sequence in U.K., potential ancestral or related sequences containing individual or combinations of amino acid changes found in the VOC202012/01 variant as identified above, and the reference genome sequence were compiled together to make the dataset for analysis. They were compiled and aligned using MAFTT (Yamada *et al*. 2016) in Jalview (Waterhouse *et al*. 2009) and then manually examined before being exported to different formats for phylogenetic tree and network analyses. Each sequence ID was renamed to contain only the GISAID accession ID number before analysis.

Phylogenetic tree analysis was conducted using MEGAX (Kumar *et al*. 2018) with maximum likelihood method, neighbor-joining method, and maximum parsimony method. A tree was constructed using each method, then bootstrapping was used to test the confidence of the phylogenetic tree from each method. For the bootstrap analysis, 500 replications were used and uniform rates among sites was assumed. The trees produced using the same method with and without bootstrapping were largely consistent with each other, so only the results from the bootstrapping analysis were presented in the supplemental results.

Phylogenetic network analysis was conducted with two different methods implemented in two different packages for comparison. The median joining method as implemented in PopART program construct network from character data and was selected based on the comparative studies of different network methods (Huson and Bryant 2006, Woolley *et al*. 2008), and it was run with epsilon set to zero. The NeighborNet method implemented in SplitsTree4 (Huson and Bryant 2006) construct network from inferred distance matrices, and the resulting network was rooted with RootedEqualAngle method.

## Data Availability

The genomic sequences of the SARS-CoV-2 used in this analysis are available from the GISAID database (https://www.gisaid.org). The sequence alignment datasets used in this study are available upon request.

## Acknowledgments

We acknowledge gratefully the authors and originating and submitting laboratories of the sequences from GISAID’s EpiCoV database on which this study is based. We also thank the GISAID data management team for answering our questions regarding sample collection date information of some sequences and making prompt corrections when needed. The research is supported by a National Science Foundation award (Award No.: 2011934).

## Author contributions

X.X. conceived and designed the research. X.X., T.L., N.G., and Z.W. performed data analysis.

X.X. wrote the paper.

## Supplemental Materials

**Supplemental Table 1.** List of the GISAID sequences used in the study.

| Accession ID (EPI_ISL ) | GISAID Clade | PANGO Lineage | Collection Date | Collection Location |
| --- | --- | --- | --- | --- |
| 402124 | L | B | 2019-12-30 | China |
| 413997 | GR | B.1.1 | 2020-02-26 | Switzerland |
| 433949 | GR | B.1.1.164 | 2020-04-12 | U.K. (England) |
| 436726 | G | B.1 | 2020-04-27 | Italy |
| 439399 | GR | B.1.1.29 | 2020-03-31 | U.K. (England) |
| 444181 | GR | B.1.1.29 | 2020-04-11 | U.K. (England) |
| 451345 | G | B.1 | 2020-01-24 | China |
| 461076 | G | B.1.8 | 2020-03-31 | Netherlands |
| 468968 | G | B.1 | 2020-03-28 | Spain |
| 480662 | GR | B.1.1.136 | 2020-06-03 | Australia |
| 481109 | G | B.1 | 2020-04-10 | Spain |
| 493480 | GR | B.1.1.29 | 2020-04-22 | U.K. (England) |
| 509505 | G | B.1 | 2020-01-25 | Australia |
| 533022 | GR | B.1.1.301 | 2020-07-15 | U.K. (England) |
| 542178 | GR | B.1.1.29 | 2020-03-18 | Italy |
| 558770 | GR | B.1.1.136 | 2020-06-08 | U.K. (England) |
| 574824 | GR | B.1.1 | 2020-03-22 | Switzerland |
| 600093 | GRY | B.1.1.7 | 2020-09-30 | U.K. (England) |
| 601443 | GRY | B.1.1.7 | 2020-09-20 | U.K. (England) |
| 631943 | GH | B.1 | 2020-05-02 | U.S.A. |
| 860053 | GR | B.1.1 | 2020-12-27 | United Arab Emirates |
| 860063 | O | B.1.1 | 2020-12-27 | United Arab Emirates |
| 860247 | G | B.1 | 2020-12-27 | Switzerland |
| 887331 | G | B.1 | 2020-03-28 | Germany |
| 896092 | GR | B.1.1 | 2021-01-29 | Switzerland |
| 896117 | G | B.1 | 2020-12-10 | Switzerland |
| 911244 | GR | B.1.1 | 2021-01-13 | Ireland |
| 953194 | GR | B.1.1 | 2021-01-03 | U.K. (England) |
| 1018090 | GR | B.1.1 | 2020-12-22 | Ghana |
| 1034923 | G | B.1.8 | 2021-02-21 | Netherlands |
| 1082422 | GR | B.1.1 | 2021-01-28 | Turkey |
| 1082424 | GR | B.1.1 | 2021-01-28 | Turkey |
| 1097167 | O | B.1.1 | 2021-02-11 | Turkey |
| 1130639 | GR | B.1.1 | 2021-01-19 | Switzerland |
| 1172307 | GR | B.1.1.220 | 2021-02-24 | U.S.A. |
| 1195103 | G | B.1.177 | 2020-11-02 | Switzerland |
| 1198083 | GRY | B.1.1 | 2021-02-02 | Sweden |
| 1210780 | GRY | B.1.1 | 2021-02-19 | Germany |
| 1211435 | O | B | 2021-01-28 | Germany |
| 1287331 | O | B.1.1 | 2021-02-17 | Germany |
| 1306807 | G | B.1 | 2021-03-13 | U.S.A. |

**Supplemental Figure 1.**
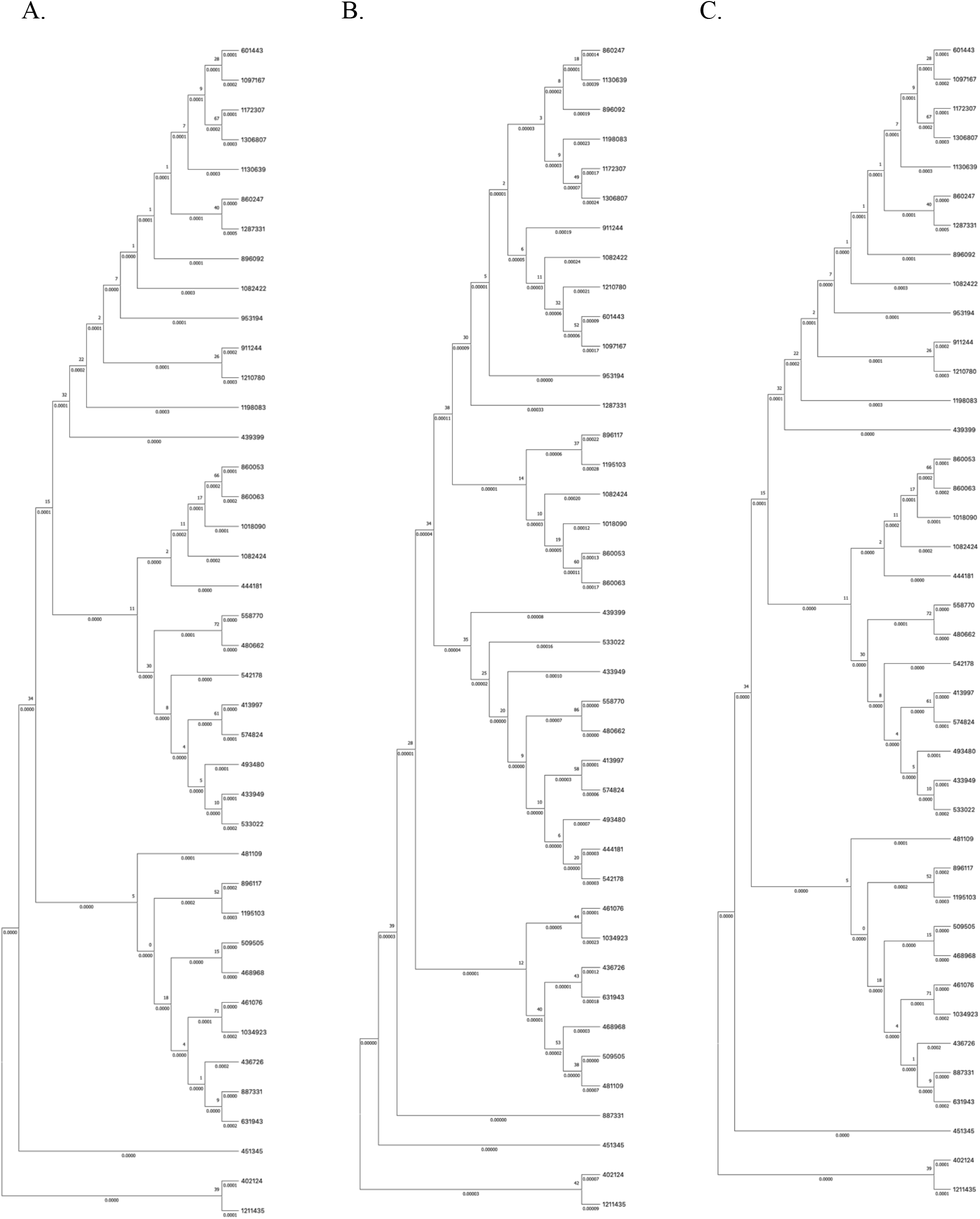
Phylogenetic tree with bootstrap statistical analyses as implemented in MEGAX. The bootstrap analysis was done with 500 replications. A. Maximum likelihood method; B. Neighbor-joining method; C. Maximum parsimony method.

**Supplemental Figure 2.**
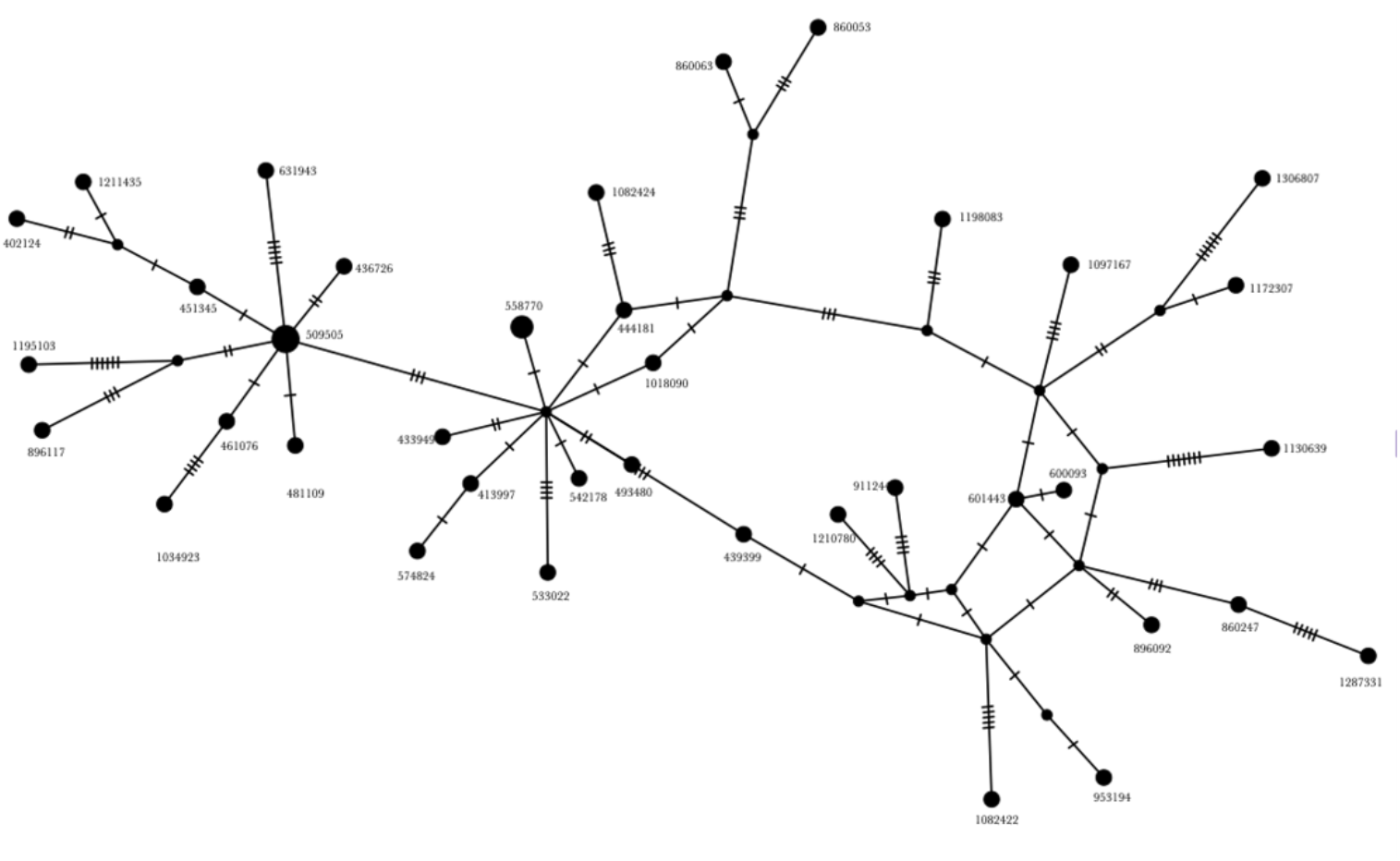
Phylogenetic network analysis with PopART and additional sequence for the VOC202012/01 variant (PANGO Lineage: B.1.1.7). Adding the additional sequence for the variant did not change the network topology or the result that the original variant sequence was a recombinant.

**Supplemental Figure 3.**
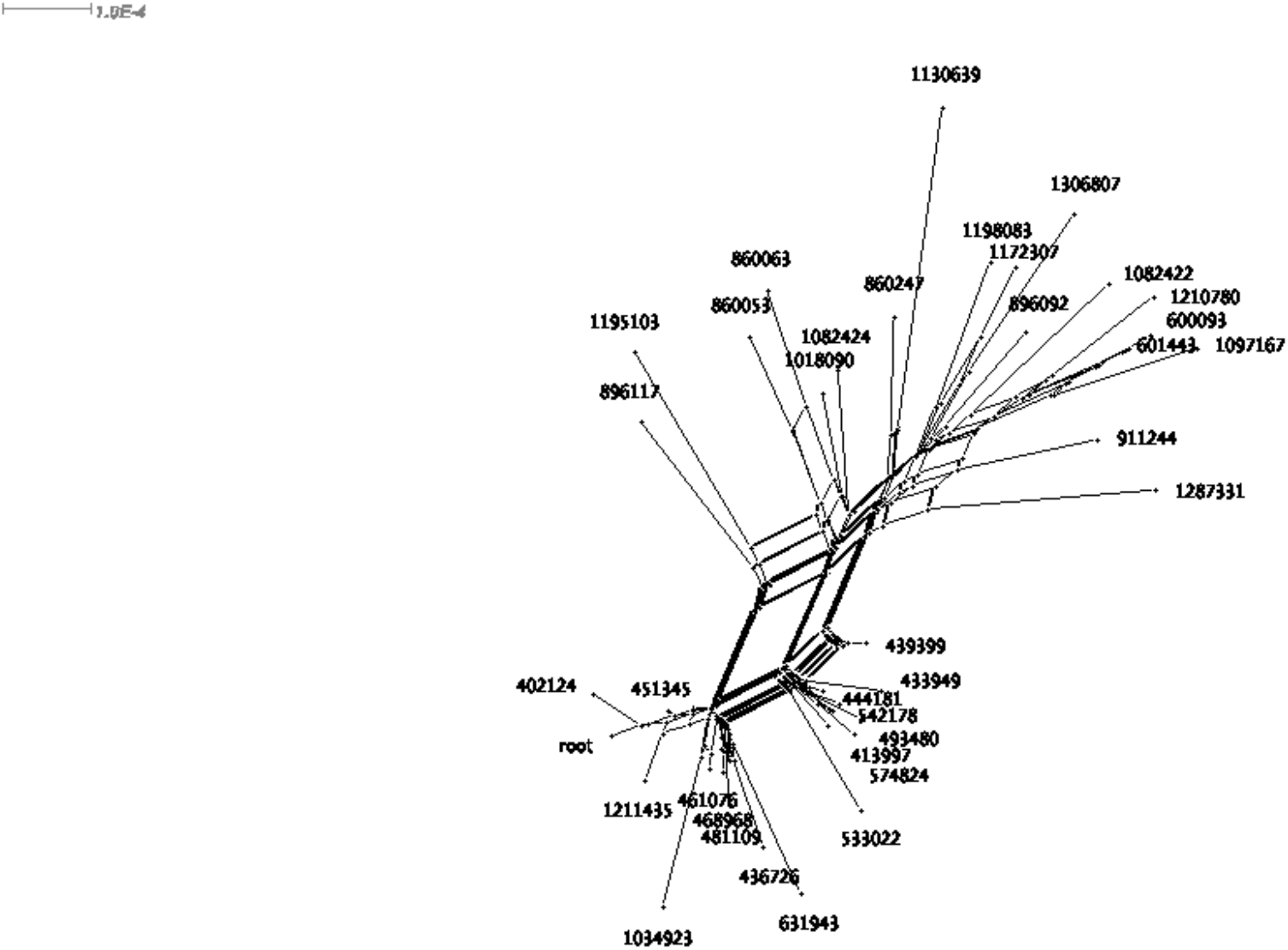
Phylogenetic network analysis with SplitsTree4 and additional sequence for the VOC202012/01 variant (PANGO Lineage: B.1.1.7). Adding the additional sequence for the variant did not change the network topology or the result that the original variant sequence was a recombinant.

